# Faecal glucocorticoid metabolite levels of intensive captive, free-range captive, and wild Tasmanian devils

**DOI:** 10.1101/2023.01.11.523346

**Authors:** Stevie Nicole Florent, Judy Clarke, Meredith J. Bashaw, Rodrigo Hamede, Menna E. Jones, Elissa Z Cameron

## Abstract

Captivity can alter the stress physiology and behaviour of an animal in both the short- and long-term through repetitive exposure to novel stressors and, subsequently, may reduce the success of conservation efforts such as translocation and reintroduction. The Tasmanian devil (*Sarcophilus harrisii*) is threatened with extinction from a fatal facial tumour disease which has led to the establishment of an insurance meta-population designed for future reintroductions of disease-free devils. The meta-population is comprised of intensive captive and free-range captive environments; however, no study has yet examined the long-term physiological implications of captivity on devils. We used non-invasive faecal glucocorticoid metabolite (FGM) monitoring to determine if there were any differences in adrenal activity between intensive captive, free-range captive, and wild devils. FGMs were not age- or sex-dependent, and we found that all population-types had similar intra-population variability and mean FGMs. In conclusion, both types of captive environment maintain stress profiles similar to wild devils.

## INTRODUCTION

All environments contain predictable and unpredictable challenges that can disrupt an individual’s homeostasis. Consequently, animals have physiological and psychological mechanisms that allow them to respond to these challenges (Deak 2007; Möstl and Palme 2002). One of these mechanisms is the stress response, which begins with hypothalamic-pituitary-adrenal (HPA) axis activation and glucocorticoid hormone (cortisol or corticosterone) release from the adrenal cortex (Axelrod and Reisine 1984; Deak 2007). At basal concentrations in normal conditions, glucocorticoids regulate metabolism and energy mobilisation for everyday activities (Busch and Hayward 2009), whilst at stress-induced concentrations, they provide an immediate energy supply and induce a cascade of adaptive physiological and behavioural responses (Deak 2007; Sapolsky *et al*. 2000).

The short-term, acute stress response can increase an individual’s survival probability by preparing the body for responding to a threat using an animal’s existing energy reserve (Busch and Hayward 2009; Martin 2009; Moberg 2000); however, when stress becomes chronic (i. e., prolonged for abnormal durations or activated repeatedly), the same adaptive mechanisms can incur a biological cost, with potentially debilitating health consequences (Linklater 2010). Chronic energy recruitment in response to stress diverts energy from core biological functions, such as immunity, bodily maintenance, and reproduction, for prolonged periods of time (Moberg 2000). This resource diversion can lead to reproductive suppression (Kirby *et al*. 2009), increased disease susceptibility (Dhabhar and McEwen 1997), severe protein loss (Teixeira *et al*. 2007), neuronal cell death (Mendl 1999), depressive-like behaviour (Brummelte and Galea 2010), and developmental failure (Chand and Lovejoy 2011).

Chronic stress-induced HPA-axis disruption can lead to a number of different individual stress profiles. Whilst negative feedback normally acts during acute stress to quickly return elevated glucocorticoids to basal levels (Dickens *et al*. 2010), under chronic stress, individuals either remain exposed to consistently elevated basal glucocorticoids (Sapolsky 1992), or their HPA axis becomes down-regulated and less responsive to stressful conditions, as a seemingly adaptive mechanism in stressful environments (Linklater *et al*. 2010). Furthermore, exposure to repetitive stressors may cause habituation, where an individual still perceives a stressor but fails to release glucocorticoids in response (Busch and Hayward 2009), or learned helplessness, where the animal releases glucocorticoids but does not alter their behaviour to avoid the stressor (Hogan *et al*. 2011). Although seemingly beneficial for stressors such as tourist exposure, habituation and learned helplessness can be maladaptive, as individuals may not adequately regulate the energy required for day-to-day activities or lose the ability to respond to future adverse stimuli, while populations may suffer reduced survivorship or low reproductive rates (Busch and Hayward 2009).

Captivity has varied, species-specific effects on glucocorticoid levels and their physiological consequences (Fischer and Romero 2019), as it can expose individuals to a number of persistent and repetitive acute stressors, including forced proximity to humans (Wells 2005) and inappropriate social conditions (Carlstead and Brown 2005). The use of non-invasive physiological monitoring, such as faecal glucocorticoid metabolite (FGM) measurements, can improve our understanding of species-specific stress profiles in captivity (Shepherdson *et al*. 2004; Watters *et al*. 2021), and is a powerful mechanism to understand how an animal perceives and responds to its environment. FGM measurements represent an aggregation of both baseline and stress-induced levels of free circulating glucocorticoids metabolised by the liver over a species-specific time period (Keay *et al*. 2006; Palme 2019; Touma and Palme 2005; von der Ohe *et al*. 2004), and are particularly advantageous as collection is simple and usually non-invasive (Millspaugh and Washburn 2004; Möstl and Palme 2002). Levels are also not influenced by minor disturbances, since the time required for circulating glucocorticoid patterns to appear in the faeces exceeds the restraint time (Palme *et al*. 2005; Sheriff *et al*. 2011); therefore, FGMs can often provide a more accurate assessment of longterm glucocorticoid levels than blood sampling (for review, see Palme 2019).

The Tasmanian devil (*Sarcophilus harrisii*) is the largest extant carnivorous marsupial (Hawkins *et al*. 2006) and is endemic to Tasmania, Australia, following extirpation from mainland Australia approximately 3000 years ago (Brüniche-Olsen *et al*. 2018). It is a medium sized, sexually dimorphic species, with males weighing between 7.5 to 14 kg and females between 6 to 9 kg, and has a life span of between five and six years in the wild (Guiler 1978) but up to eight years in captivity (W. Anthony, Devils at Cradle Wildlife Park, *pers. comm*.). The Tasmanian devil lost genetic diversity during the Last Glacial Maximum 20,000 years before present (YBP; Brüniche-Olsen *et al*. 2014) and again in the Holocene approximately 3000 YBP (Brüniche-Olsen *et al*. 2018). These historic bottleneck events, along with limited immune diversity (Morris *et al*. 2015a; Morris *et al*. 2015b), are believed to have facilitated establishment of the devil facial tumour disease (DFTD) in 1996, a transmissible cancer that has caused severe population declines (Cunningham *et al*. 2021; Hawkins *et al*. 2006; Lazenby *et al*. 2018).

As an immediate conservation action to protect against DFTD and ensure a healthy population was available for possible reintroduction, an insurance meta-population comprised of intensive captive and free-range captive isolated populations of healthy individuals was initiated (Hogg *et al*. 2015; Hogg *et al*. 2016; Hogg *et al*. 2020; Jones *et al*. 2007). The loss of the devil as the top predator of the Tasmanian ecosystem is having detrimental consequences for some native marsupial species, due to the competitive action of feral cats (Cunningham *et al*. 2020). It is therefore critical to determine the most appropriate method which ensures insurance population devils are behaviourally and physiologically suitable for reintroduction. The aim of this study was to determine if there are any differences in the mean or variability of glucocorticoid levels between intensive captive, free-range captive, and wild Tasmanian devils using non-invasive FGM measurements. No study has yet examined the long-term physiological effects of captivity on the Tasmanian devil, and with captive-to-wild translocation projects currently in execution (Grueber *et al*. 2017; Rogers *et al*. 2016), this project is particularly timely to determine if devils in captive environments have similar stress physiology as the wild population.

## METHODS

The study was conducted between 30^th^ March and 12^th^ July, 2012 in Tasmania, Australia (Table 1). Data were collected from Tasmanian devils of all age classes (one to eight years old) in intensively managed captive, free-range captive, and wild populations. Intensively managed captive populations included privately owned Wildlife Parks (*n* = 5) and research populations (*n* = 2) managed by the Save the Tasmanian Devil Program (STDP) of the Tasmanian Department of Primary Industries, Parks, Water, and Environment (DPIPWE, Table 1). Two of the Wildlife Parks are also part of the Tasmanian Devil Insurance Population, managed by the Zoo and Aquarium Association (ZAA) of Australasia and the STDP.

**Table 1.**
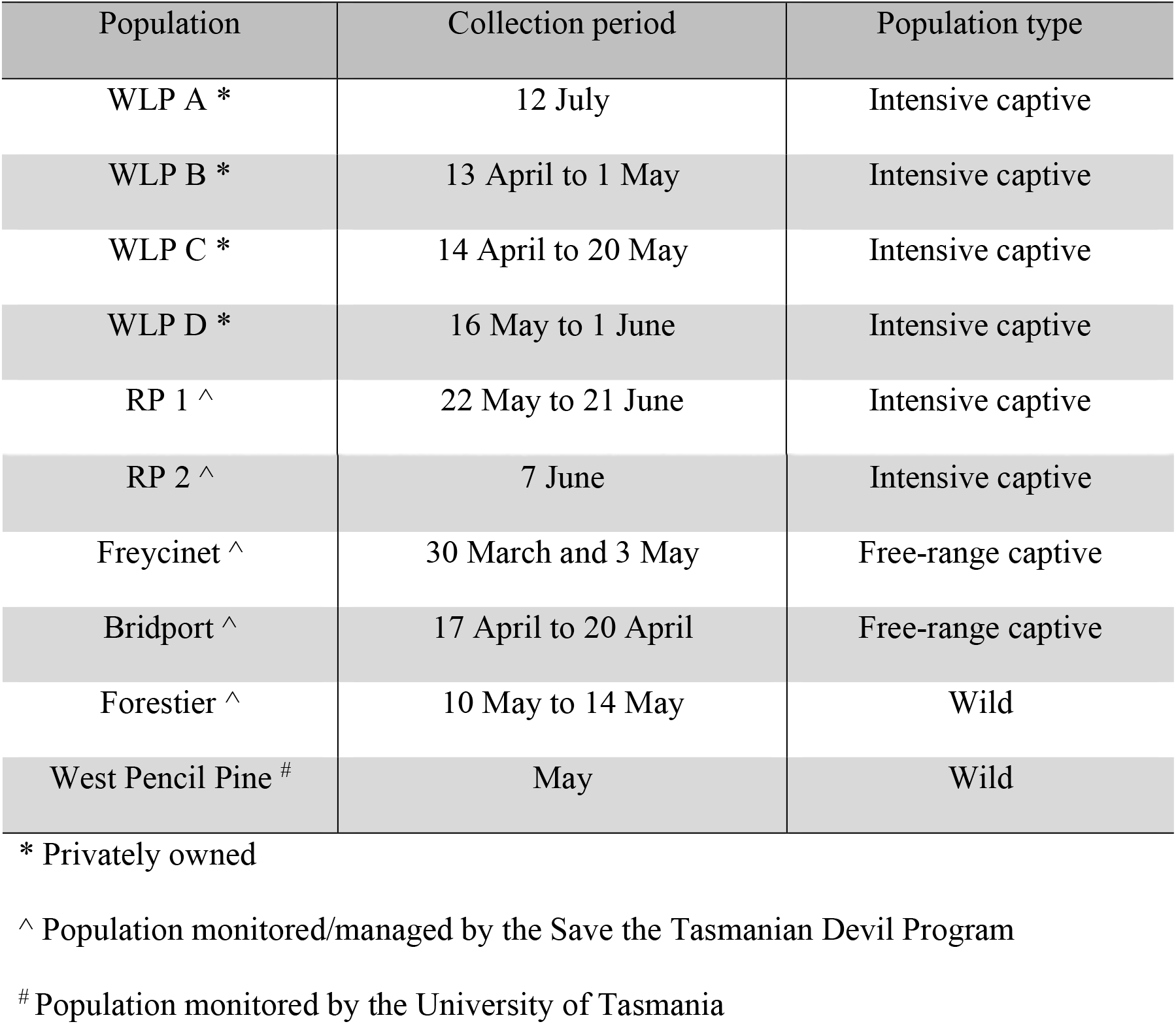
A list of the intensive captive (Wildlife Park, WLP, or research population, RP), free-range captive, and wild Tasmanian devil populations from which scat samples were collected to monitor faecal glucocorticoid metabolites, and the time period of sample collection. All samples were collected in 2012.

| Population | Collection period | Population type |
| --- | --- | --- |
| WLP A * | 12 July | Intensive captive |
| WLP B * | 13 April to 1 May | Intensive captive |
| WLP C * | 14 April to 20 May | Intensive captive |
| WLP D * | 16 May to 1 June | Intensive captive |
| RP 1 ^ | 22 May to 21 June | Intensive captive |
| RP 2 ^ | 7 June | Intensive captive |
| Freycinet ^ | 30 March and 3 May | Free-range captive |
| Bridport ^ | 17 April to 20 April | Free-range captive |
| Forestier ^ | 10 May to 14 May | Wild |
| West Pencil Pine # | May | Wild |

\* Privately owned
^ Population monitored/managed by the Save the Tasmanian Devil Program

Devils in intensive captive populations were housed in naturalistic enclosures, with dens available for shelter. Devils had access to native plants and other natural materials, including climbing structures to provide vertical complexity. Devils were kept in either same-sex or mixed-sex groups of one to ten individuals, and enclosure groupings varied during the year. Diets consisted primarily of Bennett’s wallaby (*Macropus rufogriseus*) or Tasmanian pademelon (*Thylogale billardierii*), but also other native and non-native mammals and birds (see Hesterman and Jones 2009). Fresh water was available *ad libitum*. Management regimes were not altered during the study.

Two of the three operational free-range enclosures (FREs) in Tasmania were sampled: the Bridport FRE, 20 km from the northern town of Bridport, and the Freycinet FRE on the Freycinet peninsula. Both are large-scale (22 ha) enclosures with minimal management intervention (Hogg *et al*. 2016). The FREs are double fenced with a 1.8 m high floppy-top chain mesh to reduce the probability of devils escaping, or of diseased devils entering or making physical contact with healthy FRE devils. Wallaby carcasses were provided twice a week in random locations. All FREs had *ad libitum* access to fresh water.

The two wild populations sampled were on the Forestier Peninsula in south-east Tasmania (42° 55’S, 147° 55’E) and at West Pencil Pine, 15 km west of Cradle Mountain, in north-west Tasmania (41° 31’ S, 145° 46’ E; Hamede *et al*. 2012). The Forestier Peninsula is a 160 km^2^ semi-isolated peninsula comprising private land managed for livestock production and public production forestry, and West Pencil Pine is a 25 km^2^ area located on private land managed for timber production. DFTD was prevalent in both wild populations and samples were labelled with the individual’s disease status. At the time of sampling, both populations had experienced DFTD-related decline (Hamede *et al*. 2012).

### Scat collection, storage, and processing

Entire scats were collected between the 30^th^ March and 12^th^ July, 2012 (Table 1). This period encompasses some of the mating season when females have previously been shown to have elevated corticosteroid concentrations (Keeley *et al*. 2012a; Pemberton 1990); however, time constraints required collection during this period. In intense captive populations, scats were collected by keepers during daily husbandry servicing (0900 h to 1300 h), or opportunistically during the day, and identified to the enclosure level. In the FREs and wild populations, traps were set daily (FREs: at or after 1500 h; wild populations: between 0800 h and 1600 h) and checked the following morning (FREs: from 0800 h to 1200 h; wild populations: from 0730 h onwards). Faecal samples were collected from either the trap or hessian bag used to contain the individual and were identified to an individual level. Samples on the ground at FREs judged to be < 24 h old based on their level of moisture and presence on a previous visit to that location were also collected.

The time between defecation and collection was estimated to be < 24 h for all samples. Samples were frozen at −20 °C on-site within 6 h of collection, prior to being transported and stored at −20°C until processing and analysis to prevent any further bacterial metabolism of hormone metabolites (Narayan *et al*. 2012). Storage length (≤ 5 months) was not considered a confounding factor as freezing at −20°C can reliably store faecal samples for glucocorticoid metabolite determination for up to two years (Hunt and Wasser 2003). Samples received no pre-freezing chemical treatment (Palme 2005). Scat processing and hormone extraction protocols followed those previously reported (Keeley *et al*. 2012b).

### Enzyme-immunoassay of steroids

Although cortisol is the predominant circulating glucocorticoid in devils (Weiss and Richards 1971), an assay designed to detect corticosterone better detects faecal glucocorticoid metabolites (FGM) and has previously been shown to reliably indicate stress in devils (Keeley *et al*. 2012b). Faecal sample extracts (*N* = 103) were analysed for FGM concentrations using an enzyme-immunoassay (EIA) in which FGM (diluted 1:4 in phosphate buffer) competes with horseradish-peroxidase-conjugated corticosterone (1:20,000) to bind with a polyclonal antibody (CJM06, 1:12,500) following Keeley *et al*. (2012b) and Narayan *et al*. (2012).

Sample binding was compared to corticosterone standards (4.88 – 2500 pg per 50 μL; Steraloids Inc., USA) to determine FGM concentrations. In this study, the percent binding for serially diluted pooled faecal samples (1:1 to 1:128) from each population type (wild, *n* = 9; FRE, *n* = 10; intensive captive, *n* = 10) were parallel to the corticosterone standard preparations, indicating that the assay was effective and comparable for all three populations. Any samples which bound outside the linear part of the parallelism curve were re-run at a lower dilution (1:2). Duplicates with a coefficient of variation of >15% were also re-run. Hormone concentrations were expressed as nanograms per gram (ng/g) of dried faecal material (DFM).

### Statistical analysis

All statistical analyses were performed using SPSS (versions 19.0.0.1 and 28.0.0.0, SPSS, Inc., 2021, IBM©). All data are presented as mean ± 1 SEM unless otherwise stated. For all statistical tests, significance was accepted at *p* ≤ 0.05. Homogeneity of variances and normality of data distribution were assessed by visual inspection of: 1) group standard deviations versus group means plots; and 2) the normal plots of residuals versus predicted residual values, respectively. All post-hoc analyses were conducted using Tukey’s HSD test. FGM concentrations were natural log transformed prior to analysis to meet normality and homoscedasticity assumptions of a parametric test (Hamilton 1992), and all mentions of FGM refer to the natural log transformed values unless otherwise stated. An independent samples t-test found sample age prior to freezing did not influence FGM concentrations (0 to 11 h old, n = 28, 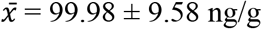 dried faecal material; 12 to 24 h old, n = 28, 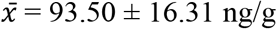 dried faecal material; *t* _53_ = 1.06, *p* = 0.30), and thus it was not included in future analyses (Millspaugh and Washburn 2004).

Univariate General Linear Models (GLM) were used to determine if an individual’s age and/or sex influenced FGM measurements (response variable). Only samples from individuals or intensive-captive enclosure groupings of known age and sex were included in this analysis. The independent variables were the population type (i.e., intensive captive, *n* = 31; free-range captive, *n* = 17; or wild, *n* = 7), the age class (‘subadult’, ≤ 2 years old, *n* = 29; ‘adult’, >2 years old, *n* = 26), and sex (female, *n* = 28; male, *n* = 27). Reproductive status was not incorporated into analyses, as the sample size of individuals with known reproductive status was too low. A second univariate GLM was applied to determine if enclosure sex composition in Wildlife Parks influenced FGM (male-only, *n* = 8; female-only, *n* = 12; mixed sex groups, *n* = 25). Following this, a final univariate GLM was used to test for variance in FGM measurements (response variable) in relation to the population type (independent variable; intensive captive, *n* = 70; free-range captive, *n* = 23; wild, *n* = 10).

A Levene’s test of equality of error variances tested if FGM error variance was equal across groups. FGM measurements were further analysed by categorical subdivision into ‘high’ (>100 ng/g DFM, *n* = 37) and ‘low’ (≤100 ng/g DFM, *n* = 66) levels in each population type, and a Pearson’s chi-square test was used to determine if there was a significant difference amongst population types. The individual Wildlife Park or free-range enclosure the sample was from was not considered, as we were more interested in testing for overall differences between population types. Lastly, a univariate GLM was run to determine if an individual’s disease status influenced FGMs for the six wild samples that had this information (not diseased, *n* = 4; diseased, *n* = 2).

## RESULTS

FGM levels were similar between subadult (≤ 2 years old) and adult devils (> 2 years old; subadult, *n* = 29, 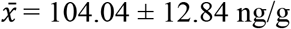 DFM; adults, *n* = 26, 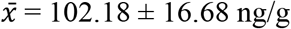 DFM; Table 2). There was no three-way interaction between population type and an individual’s age class and sex, nor a two-way interaction between age class and sex (Table 2).

**Table 2.**
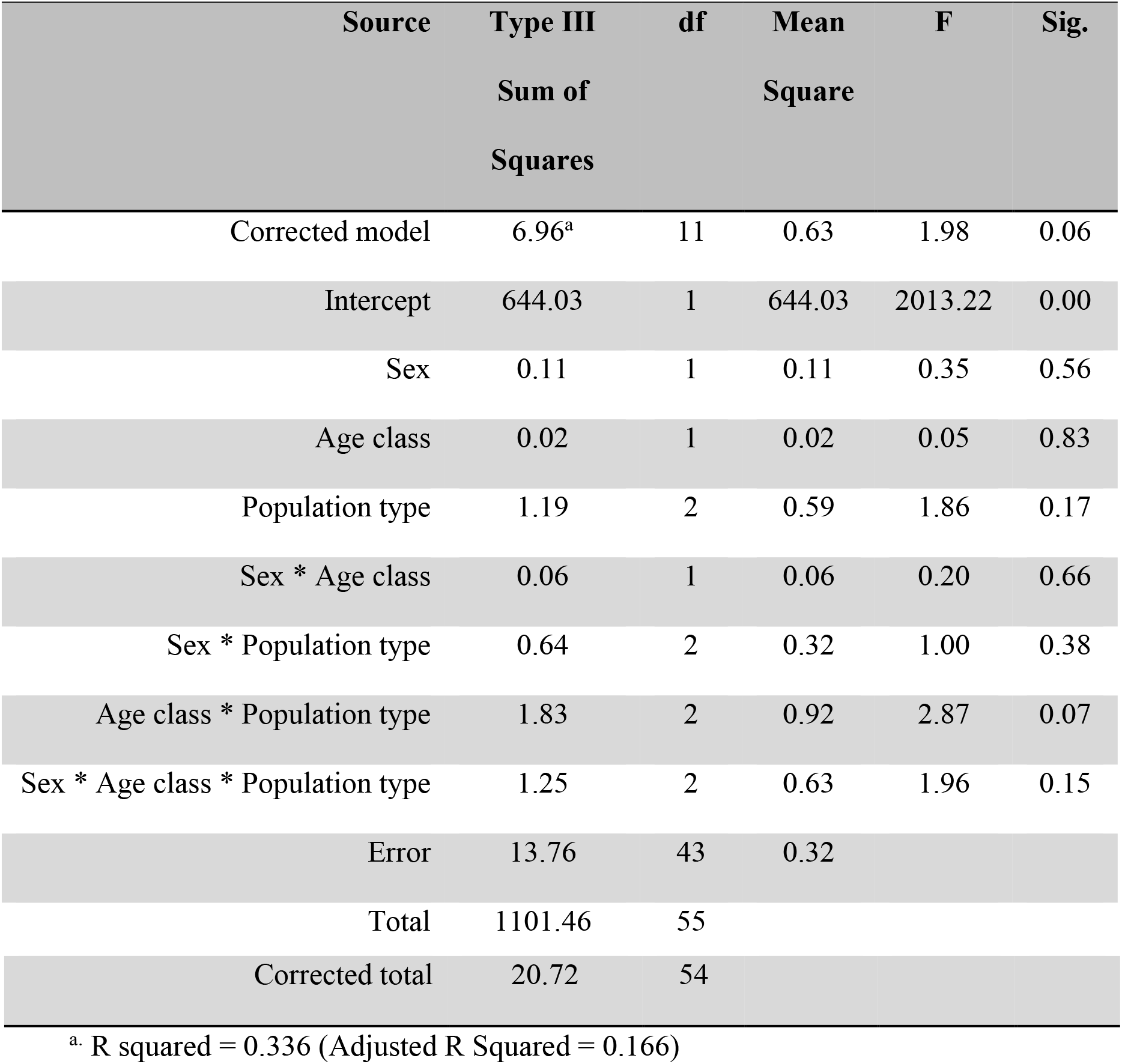
Univariate general linear model examining the potential effects of age class (subadult or adult), sex, and population type (intensive captive, free-range captive (FRE), and wild) on log-transformed faecal glucocorticoid metabolite (FGM) concentrations of Tasmanian devils.

| Source | Type III<br>Sum of<br>Squares | df | Mean<br>Square | F | Sig. |
| --- | --- | --- | --- | --- | --- |
| Corrected model | 6.96<sup>a</sup> | 11 | 0.63 | 1.98 | 0.06 |
| Intercept | 644.03 | 1 | 644.03 | 2013.22 | 0.00 |
| Sex | 0.11 | 1 | 0.11 | 0.35 | 0.56 |
| Age class | 0.02 | 1 | 0.02 | 0.05 | 0.83 |
| Population type | 1.19 | 2 | 0.59 | 1.86 | 0.17 |
| Sex * Age class | 0.06 | 1 | 0.06 | 0.20 | 0.66 |
| Sex * Population type | 0.64 | 2 | 0.32 | 1.00 | 0.38 |
| Age class * Population type | 1.83 | 2 | 0.92 | 2.87 | 0.07 |
| Sex * Age class * Population type | 1.25 | 2 | 0.63 | 1.96 | 0.15 |
| Error | 13.76 | 43 | 0.32 |  |  |
| Total | 1101.46 | 55 |  |  |  |

|  |  |  |
| --- | --- | --- |
| Corrected total | 20.72 | 54 |

<sup>a</sup>. R squared = 0.336 (Adjusted R Squared = 0.166)

A trend for a two-way interaction between age class and population type (GLM, *F*_2,50_ = 2.87, *p* = 0.07) was found, with FRE adults (*n* = 6) displaying visually lower FGMs than wild (*n* = 2) and intensive captive (*n* = 18) adults; however, as the result was insignificant and sample sizes were low, age was not included in further analyses, and all non-age specific samples were used (Fig. 1).

**Fig. 1.**
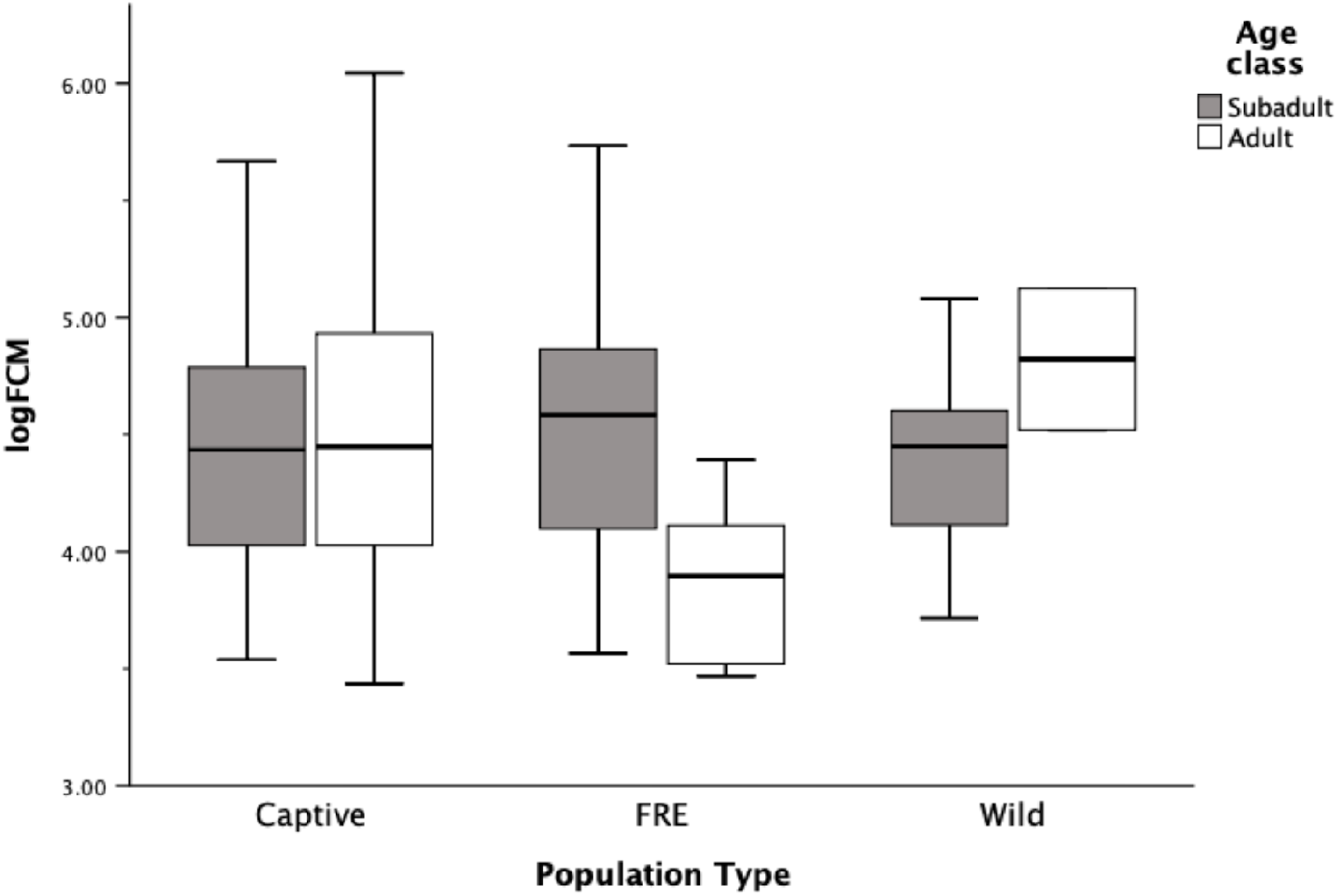
Log-transformed faecal glucocorticoid metabolite (FGM) concentrations (nanograms per gram (ng/g) dried faecal material) of adult (> 2 years old, *n* = 26) and subadult (≤ 2 years old, *n* = 29) Tasmanian devils in intensive captive (captive), FRE, and wild populations in Tasmania, Australia (captive: adults, *n* = 18, subadults, *n* = 13; FREs: adults, *n* = 6, subadults, *n* = 11; wild: adults *n* = 2, subadults *n* = 5).

An individual’s sex did not influence FGM levels (females, *n* = 28, 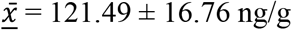 DFM; males, *n* = 27, 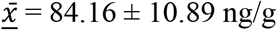 DFM; GLM, *F_1,53_* = 0.35, *p* = 0.56, Fig. 2A), and there was no interaction between sex and population type (GLM, *F*_2,50_ = 01.00, *p* = 0.38, Table 2), so sex was not included in further analyses, and all samples including non-sex specific samples were used. Despite low sample sizes, the known seasonal pattern of female glucocorticoids was observed (Fig 2B), with a peak in April followed by a decline to a low in June (Fig. 3, Keeley *et al*. 2012a; Pemberton 1990).

**Fig. 2.**
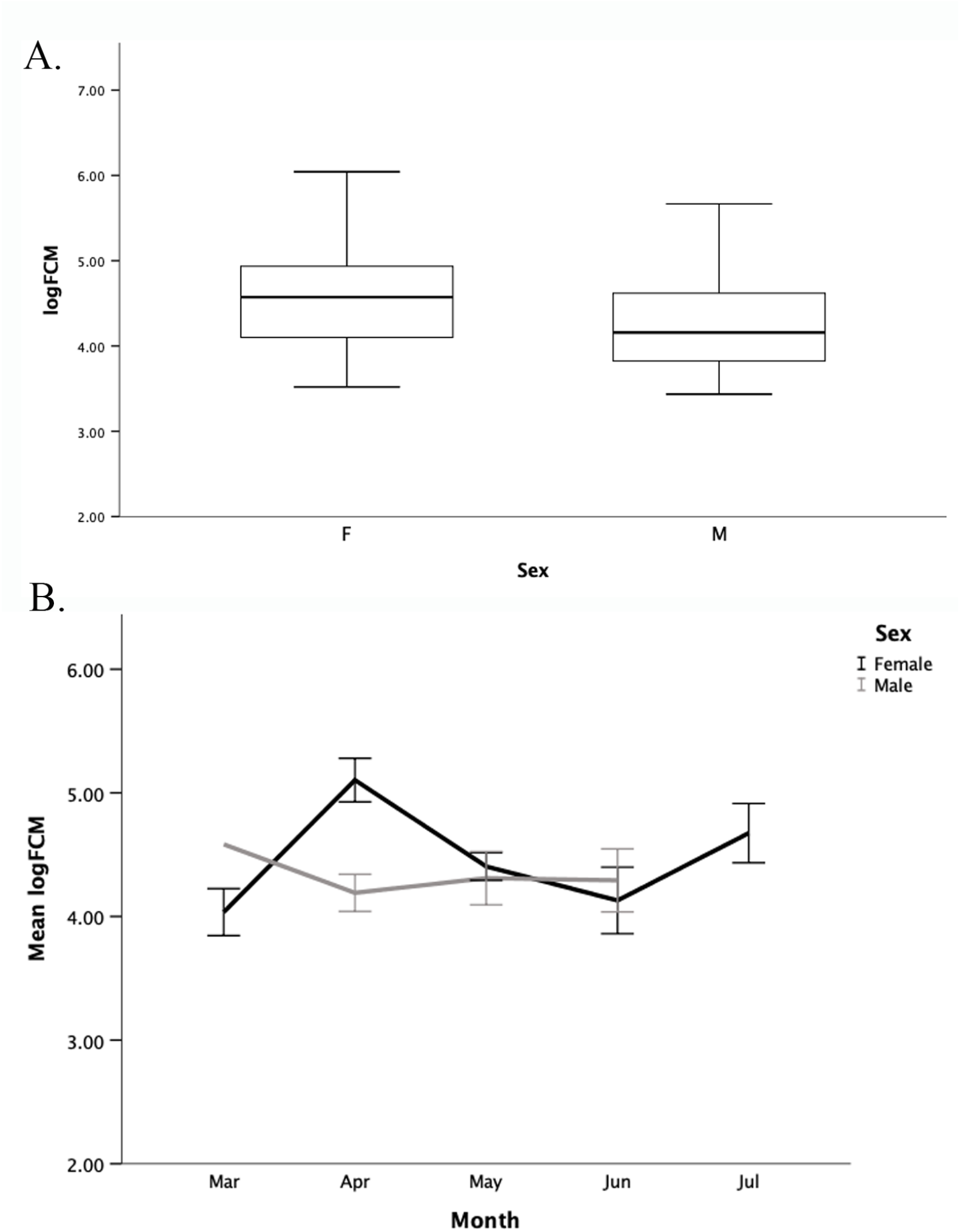
(A) Log-transformed faecal glucocorticoid metabolite (FGM) concentrations (nanograms per gram (ng/g) dried faecal material) for female (*n* = 28, F) and male (*n* = 27, M) Tasmanian devils from populations in Tasmania, Australia. (B) Mean log-transformed FGM concentrations for female (black line) and male (light grey line) Tasmanian devils from March to July, 2012. Only one male-identified sample was collected in March, and no male-identified samples were collected in July. Data presented as mean ± SEM.

**Fig. 3.**
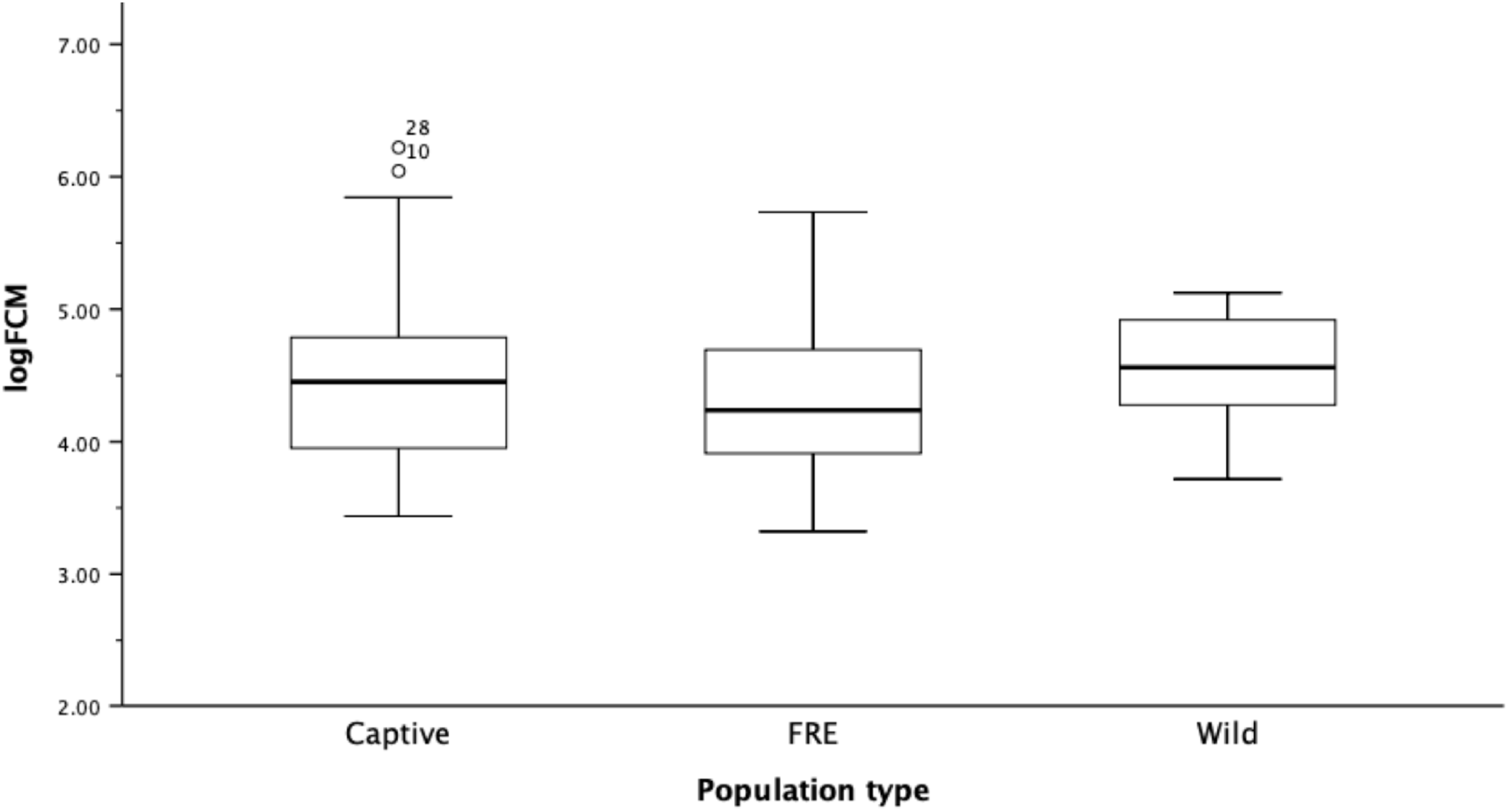
Log-transformed faecal glucocorticoid metabolite (FGM) concentrations (nanograms per gram (ng/g) dried faecal material) in intensive captive (*n* = 70), free-range captive (FRE, *n* = 23), and wild (*n* = 10) Tasmanian devils from populations in Tasmania, Australia. Outlier #10 an adult female housed separately with young, whilst #28 was a sample from a mixed-sex intensive captive enclosure.

From the second univariate GLM, enclosure sex composition in the intensive captive population did not influence FGM concentrations (female-only, *n* = 12, 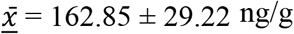 DFM; male-only, *n* = 8, 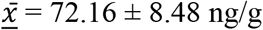 DFM; mixed-sex, *n* = 25, 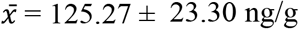 DFM; GLM, *F*_2,42_ = 1.96, *p* = 0.16), nor did the number of animals in the enclosure (GLM, *F*_5,39_ = 1.53, *p* = 0.22).

Population type did not influence FGM concentrations (Table 3, Fig. 3). Furthermore, there was no significant difference in FGM concentrations between population types when separated categorically into ‘high’ (>100 ng/g DFM, *n* = 37) and ‘low’ (≤100 ng/g DFM, *n* = 66) FGM levels (Pearson’s chi-squared test, χ^2^ = 0.42, *df* = 1, *p* > 0.20). Interestingly, the inter-individual variation present in each population was similar (Levene’s test, *F*_2,100_ = 0.83, *p* = 0.44). Intensive captive, free-range captive, and wild populations thus showed equivalent means and variance in FGM concentrations (intensive captive, *n* = 70, 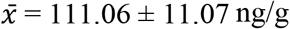 DFM; free-range captive, *n* = 23, 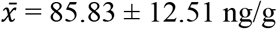 DFM; wild, *n* = 10, 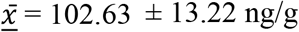 DFM).

**Table 3.**
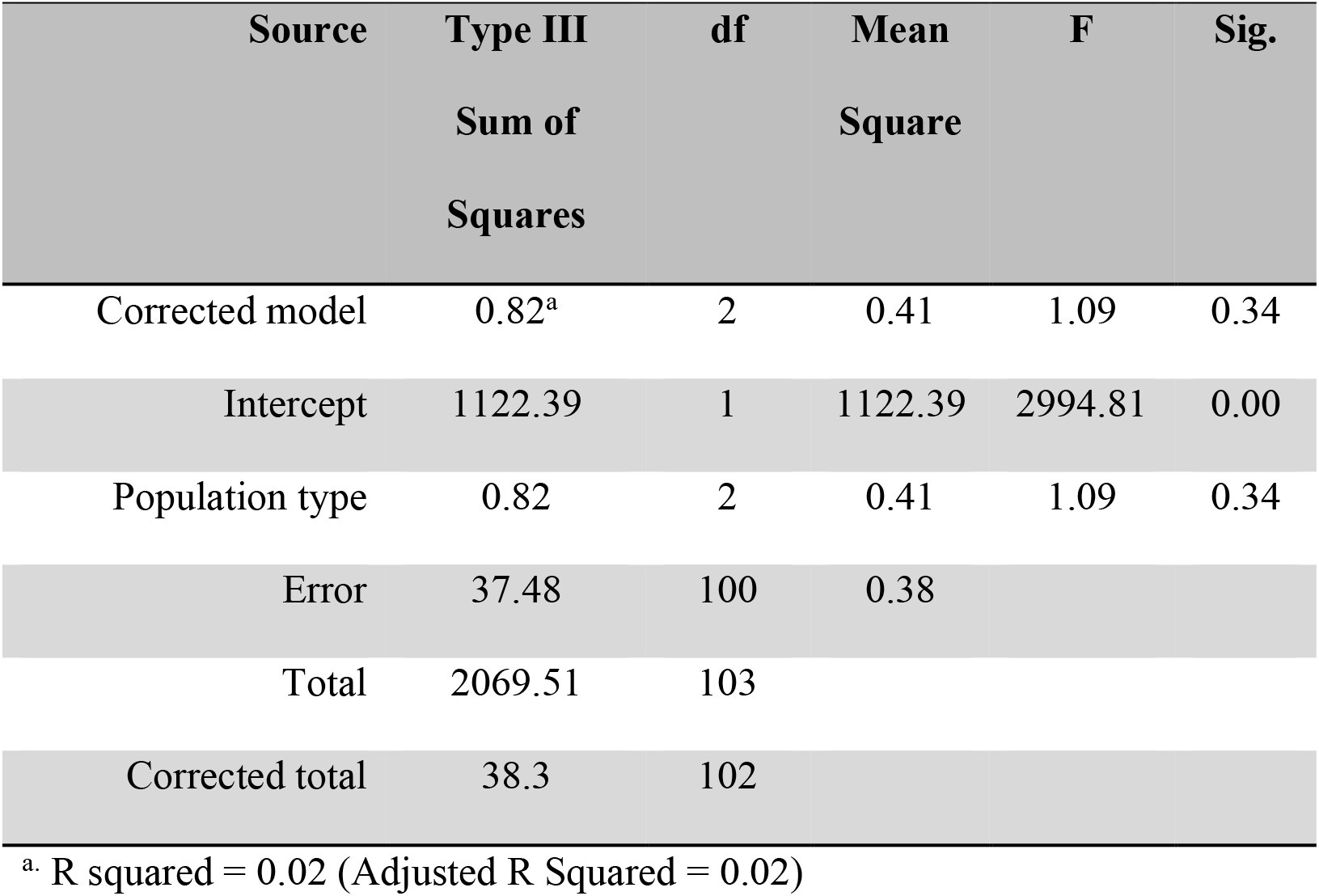
Univariate general linear model examining the potential effects of population type (intensive captive, free-range captive (FRE), and wild) on log-transformed faecal glucocorticoid metabolite (FGM) concentrations of Tasmanian devils.

Likely due to small sample sizes, there was no evidence that individual disease status influenced FGM concentrations (diseased, *n* = 2, 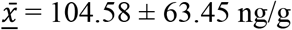 DFM; not diseased, *n* = 4, 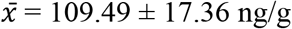 DFM; GLM, *F*_1,4_ = 0.25, *p* = 0.64).

## DISCUSSION

We found no evidence of either suppressed or elevated stress in intensive captive and free-range captive Tasmanian devils compared to the wild population, and comparable variability in FGM levels amongst and within populations. This intra-population variability is similar to that previously reported for captive female devils (Keeley *et al*. 2012b). As FGM concentrations provide an average glucocorticoid measure of the previous 24-hours (Keeley *et al*. 2012b), our results may reflect both baseline glucocorticoids and a response to acute environmental stressors, such as interspecific interactions, trapping, or human presence over this period. Although devils would experience different stressors in captivity than in the wild, the similar intra-population variability suggests the impact of these stressors is of equal magnitude. Our results of similar stress profiles between management styles are also in line with Rogers *et al*. (2016), who found the style of captive management doesn’t influence the success of Tasmanian devils released into the wild.

Variability is essential for adequate stress functioning (Sapolsky 1994), and the similar intrapopulation variability presented here suggests that captive devils retain their ability to elicit a stress response (Shepherdson *et al*. 2004). In contrast, low variability would suggest habituation (Busch and Hayward 2009), which although can seem beneficial by allowing individuals to maintain good body condition in captivity, may mean individuals have lost the ability to produce a stress-mediated response to cope with future stimuli, and consequently may suffer lower survival (Cabezas *et al*. 2007; Zidon *et al*. 2009). For example, the number of generations a devil has been in captivity is associated with vulnerability to vehicle strike which may reflect naivety or habituation (Grueber *et al*. 2017). However, the similar means and variability in FGMs in this and previous studies (Keeley *et al*. 2012b), indicates that captive devils could be experiencing normal basal levels of stress and retain their ability to elicit a stress response.

An alternative explanation for the similar inter-population FGM levels is that devils could be experiencing equivalent moderately elevated basal or ‘intermediate’ levels of stress (Busch and Hayward 2009; Cabezas *et al*. 2007; Meylan *et al*. 2012; Pravosudov 2003). Captivity exposes individuals to a multitude of novel stimuli, such as the presence of people and a greater rate of social interactions than wild individuals would experience. Intermediate stress levels, particularly when within the normal range of baseline glucocorticoids routinely experienced by wild conspecifics, can promote an adequate stress response to environmental agitations and can, therefore, enhance survival (Meylan *et al*. 2012; Pravosudov 2003). For example, intermediate cortisol levels enhance associative and spatial learning in captive juvenile Belding’s ground squirrels (*Spermophilus beldingi;* Mateo 2006). In addition, the primary role of glucocorticoids in normal conditions is to regulate energy through acquisition, deposition, and mobilisation (Busch and Hayward 2009). If an individual has suppressed glucocorticoids, they may not be able to mobilise the energy required for everyday activities. Conversely, elevated glucocorticoids can lead to fat deposition and skeletal muscle catabolism (Shepherdson *et al*. 2004). Intermediate levels of basal glucocorticoids can therefore have many beneficial outcomes, and this could be what devils are experiencing in the populations sampled in our study.

Along with comparable inter- and intra-population variability, there was also no significant difference in FGM mean levels between adult and subadult devils. FRE adults had a near-significant trend for lower FGM levels than wild or intensive-captive adults; however, we are conscious of making any inferences given the low number of samples with known age and sex status. There was also no influence of sex on FGM levels. This was surprising, as females of many species have higher basal circulating glucocorticoids than males, including rats and mice (Chisari *et al*. 1995), Siberian hamsters (*Phodopus sungorus;* Bilbo and Nelson 2003), and red-backed voles (*Clethrionomys gapperi;* Kramer and Sothern 2001), possibly due to physiological HPA and hypothalamic-pituitary-gonadal axis cross-reactivity (Handa *et al*. 1994). Although male devil reproduction is not seasonally restricted (Keeley *et al*. 2012a; Pemberton 1990), female devil free corticosteroids steadily increase from January to April, when a peak in births occur. They then decrease to a low in June before a secondary peak in October when young devils begin to emerge from dens, suggesting an influence of reproductive events on corticosteroid levels (Hesterman *et al*. 2008; Pemberton 1990). The samples in this study were collected during and following the mating season and, although restricted by sample size, we witnessed a similar trend in seasonal variation, with a peak in female FGM levels in April, followed by a decline to a low in June. Secondly, an outlier in the population graph (#10) was a female housed separately in intensive-captive population with young, possibly experiencing elevated stress due to the lactation demands of pups.

In conclusion, we found no physiological indicators of either elevated or suppressed levels of stress in intensive captive or free-range captive devils when compared to the wild population. We also show that individual devils have variable basal glucocorticoid levels regardless of population type, supporting the conclusions of Keeley *et al*. (2012b). Future longitudinal physiological studies are required to understand the devil stress response, including the metabolites formed, the primary excretion route, and the effects of various stressors on their physiological state (Teixeira *et al*. 2007). These longitudinal studies would help differentiate between the two possible nonexclusive explanations of the FGM results in our study, that is: 1) that devils are experiencing intermediate basal levels of stress; and 2) that devils retain their ability to respond to acute stressors. Additionally, future integration of behavioural and physiological indices of stress would provide a more complete picture of how an individual perceives its environment (Shepherdson *et al*. 2004; Watters *et al*. 2021), which can improve the success of active conservation management techniques (Tarszisz *et al*. 2014). The rapid and extensive decline of the devil will require active management for future decades (Jones *et al*. 2007), and consideration of individual stress profiles is vital to ensure captive devils remain physiologically and behaviourally functional representations of the species. Selecting individuals for translocation and reintroduction projects based on knowledge of their behavioural and physiological phenotype may be important for increasing the success of these future efforts.

## ACKNOWLEDGEMENTS

The authors would like to thank all owners and staff at the following Wildlife Parks for their assistance in this project: Bonorong Wildlife Sanctuary, Zoo Doo Wildlife Park, Devils at Cradle, East Coast Nature World, and Trowunna Wildlife Park. Furthermore, the authors would like to thank the Save the Tasmanian Devil Program and the Zoo and Aquarium Association of Australasia. The study was approved by the University of Tasmania Animal Ethics Committee (A12367 and A11696) and field collection approved by the Tasmanian Government under Scientific Permit TFA12200. The study was approved by the Zoo and Aquarium Association of Australasia and the Save the Tasmanian Devil Program, part of the Tasmanian Department of Primary Industries, Water, and Environment.

## Notes

### Competing Interest Statement

The authors have declared no competing interest.

